## Supplemental Information for "The evolution of ovary-biased gene expression in Hawaiian *Drosophila*"

### The evolution of ovary-biased gene expression in Hawaiian *Drosophila* - Supplementary methods and tables

### 1 Supplementary figures

#### 1.1 Background

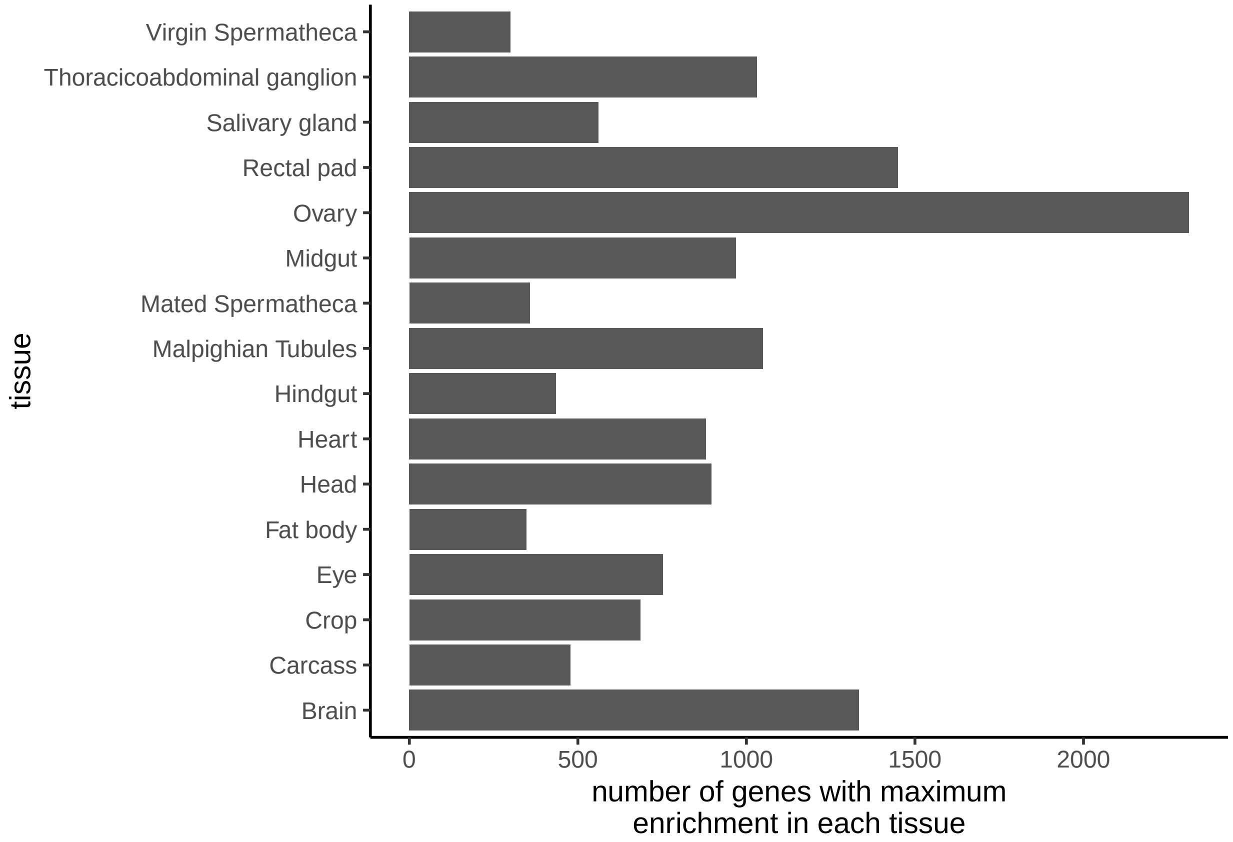

Figure S1: **Comparing across *Drosophila melanogaster* female tissues in the FlyAtlas2 dataset[1], more genes show highest enrichment in the ovary than any other tissue.**

#### 1.2 Analysis pipeline

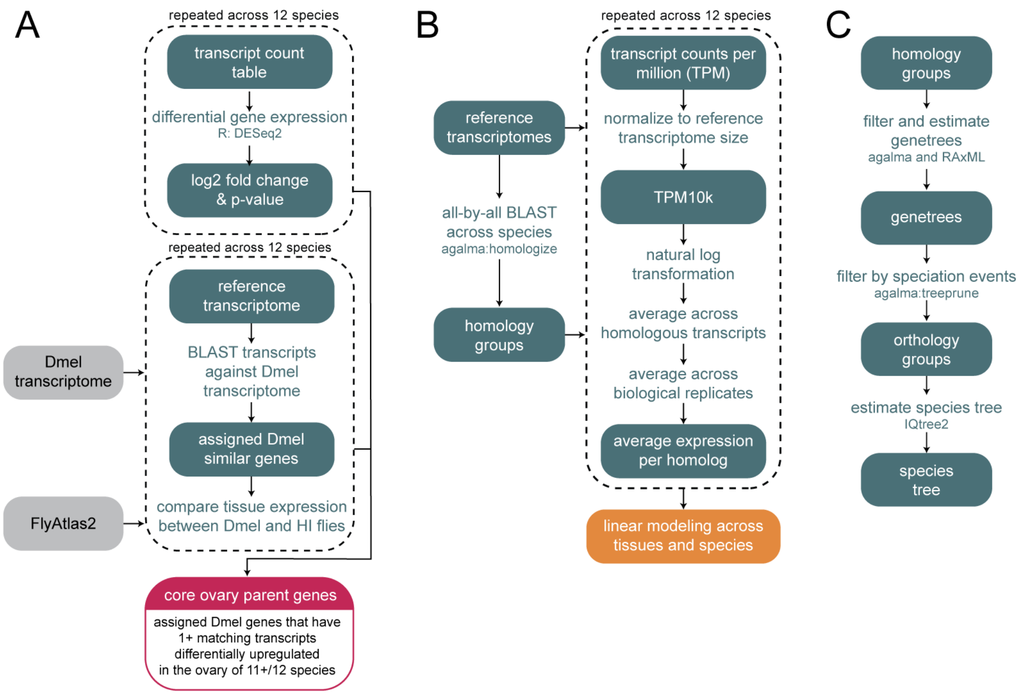

Figure S2: **Analysis pipeline for (A) identifying core ovary genes, (B) linear modeling of expression, and (C) estimating the species level phylogeny. Dmel = *Drosophila melanogster*. HI flies = Hawaiian Drosophilidae.**

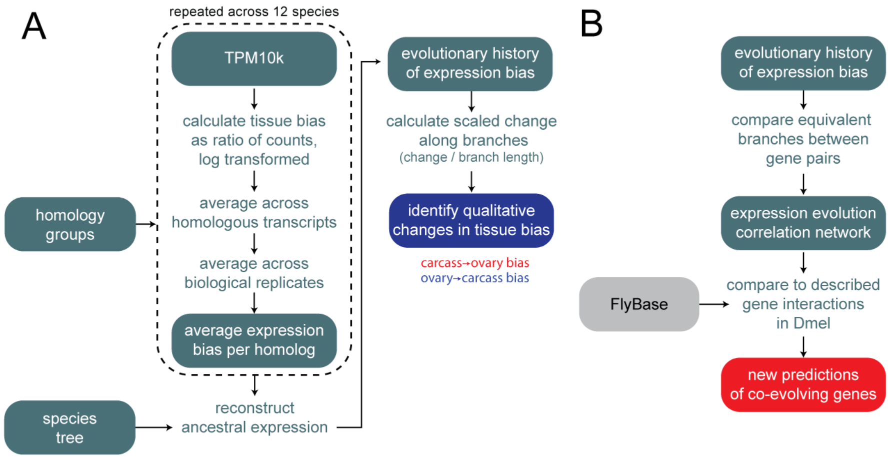

Figure S3: **Analysis pipeline for (A) identifying evolutionary changes in expression bias, and (B) estimating correlation of expression evolution between genes.**

#### 1.3 Differential gene expression

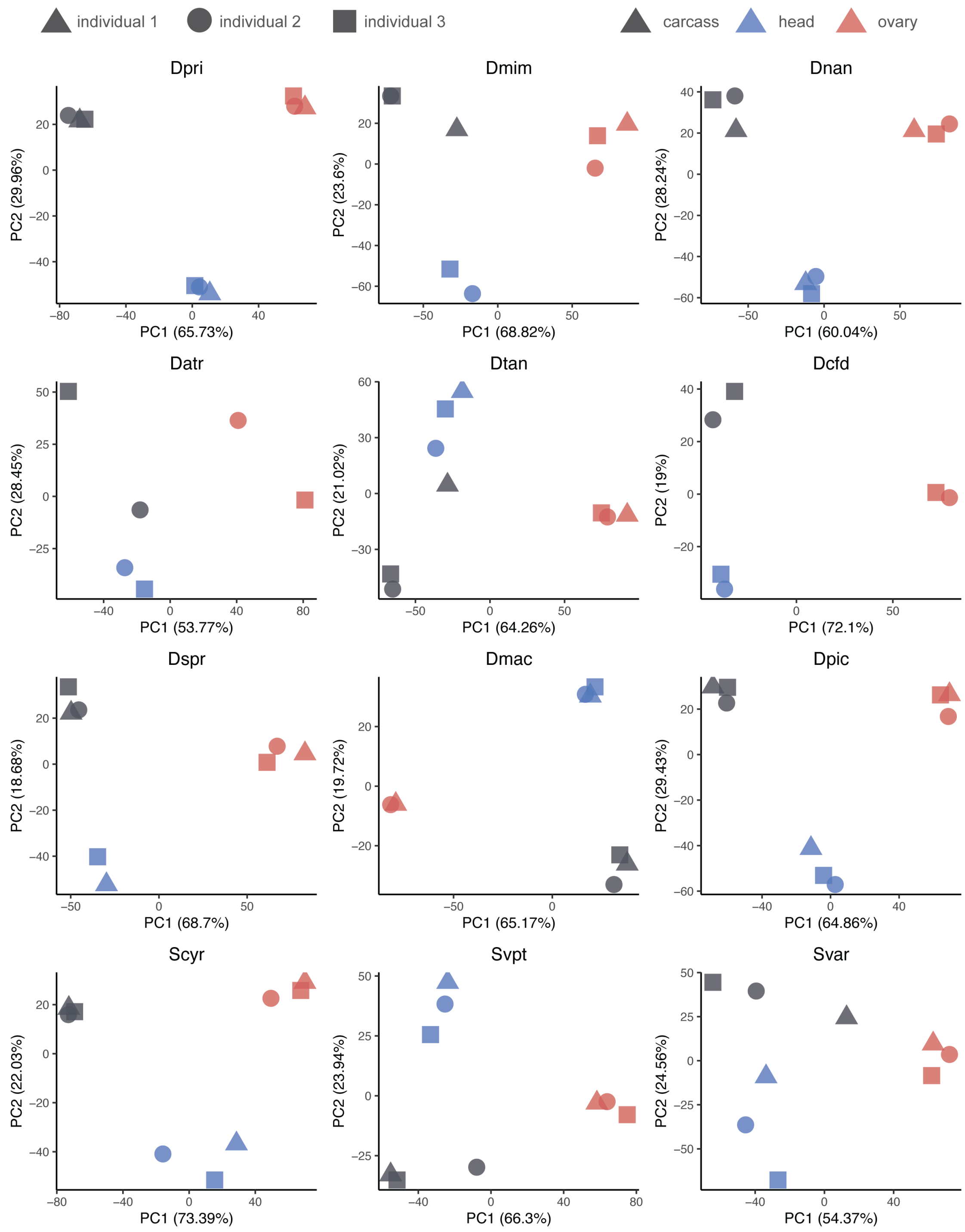

Figure S4: **Principal component analysis within species.** Species abbreviations correspond to names shown in Fig. 1B.

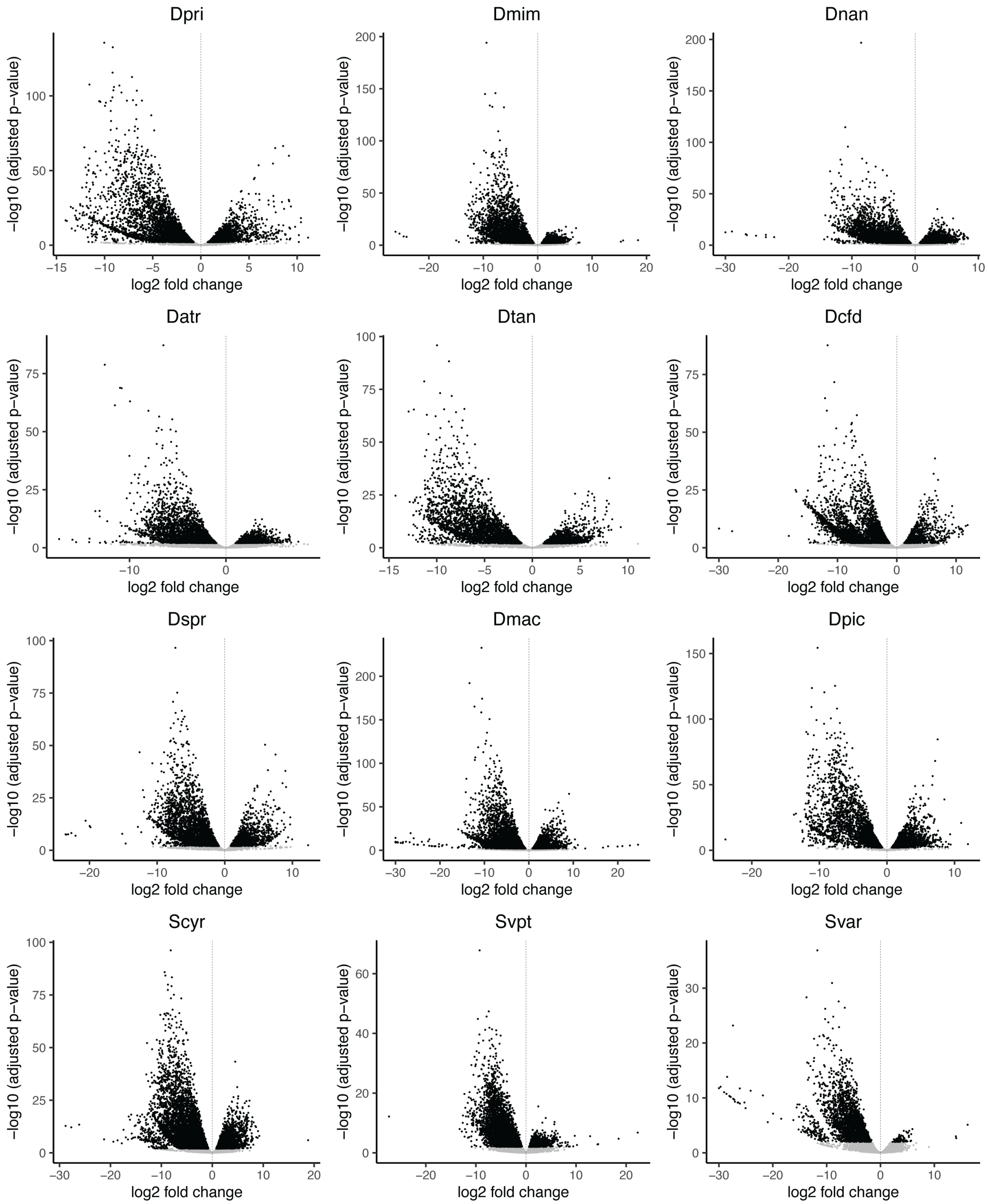

Figure S5: **Differential gene expression volcanos plot comparing ovary and carcass across species.** Species abbreviations correspond to names shown in Fig. 1B. Positive fold changes (points on the right hand side of the volcano) indicate higher expression in the ovary than in the carcass.

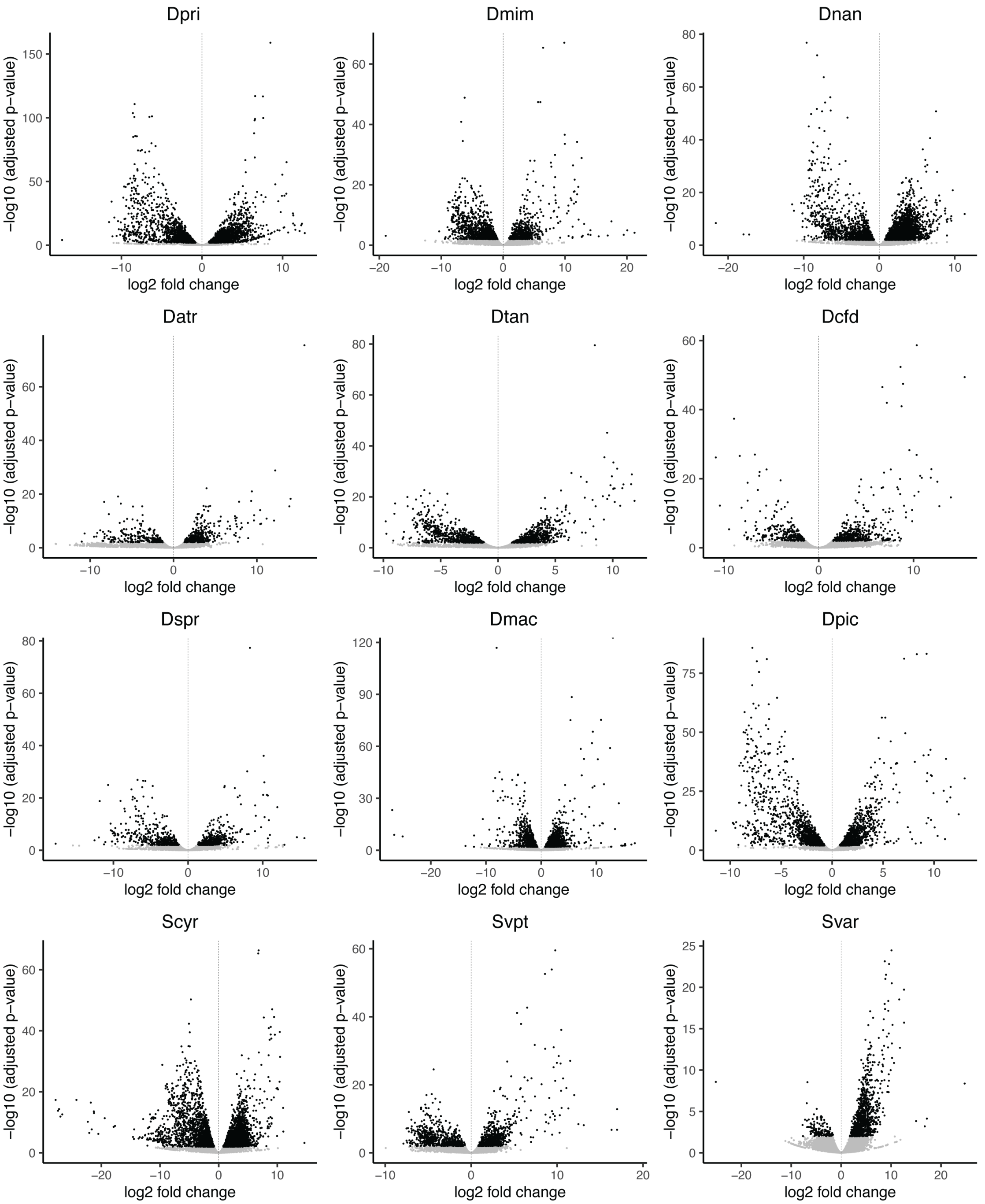

Figure S6: **Differential gene expression volcanos plot comparing head and carcass across species.** Species abbreviations correspond to names shown in Fig. 1B. Positive fold changes (points on the right hand side of the volcano) indicate higher expression in the head than in the carcass.

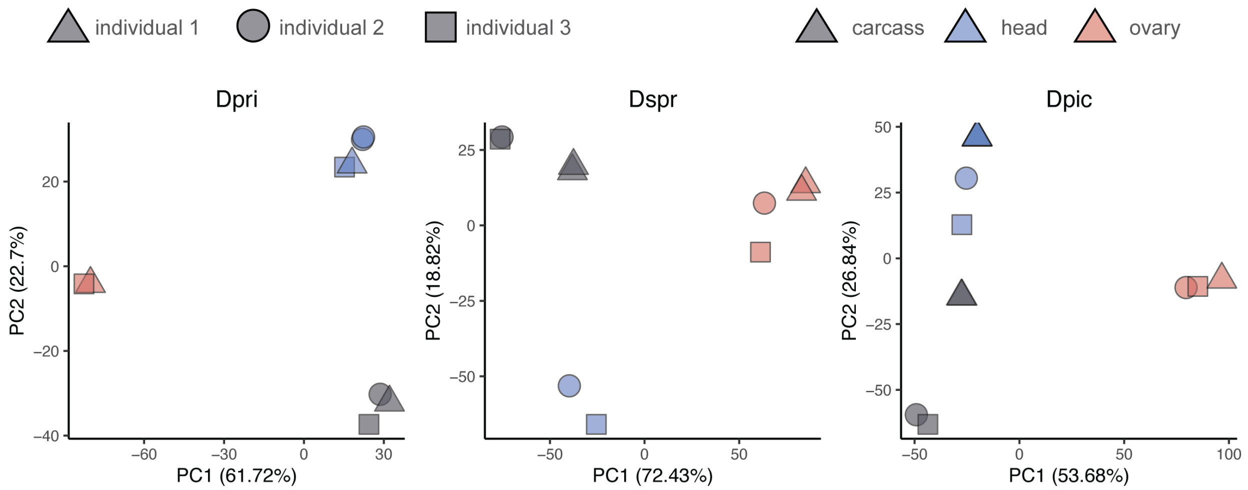

Figure S7: **Principal component analysis showing resequenced libraries.** Species abbreviations correspond to names shown in Fig. 1B.

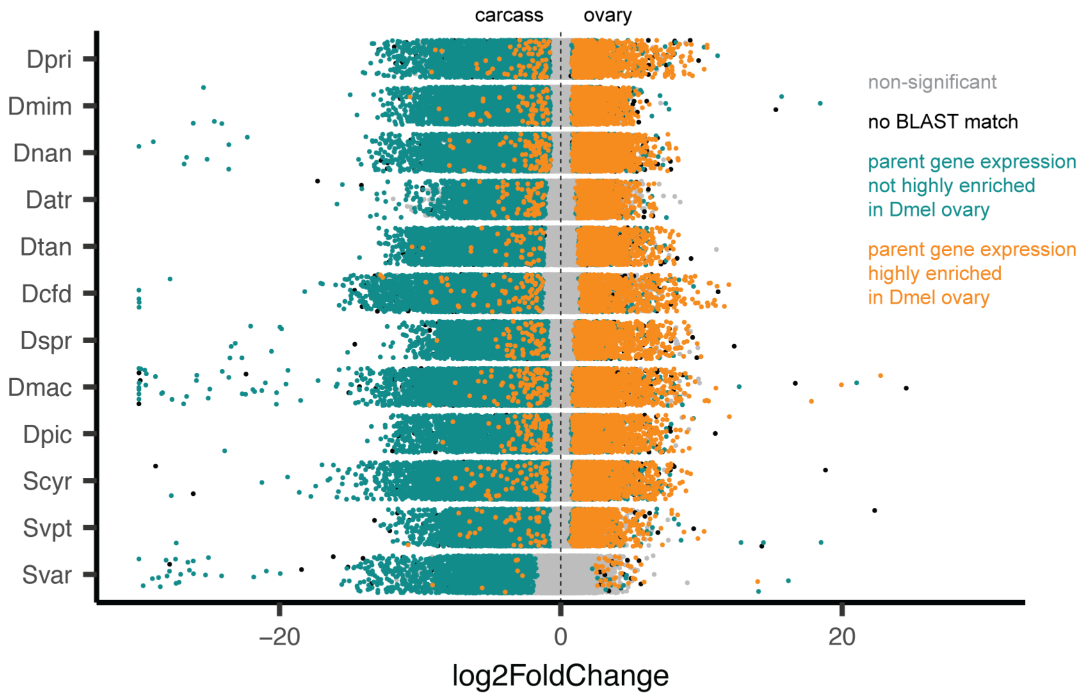

Figure S8: **Differential gene expression analysis across species, colored by *D. melanogaster* expression enrichment.** Points in cyan indicate transcripts that match *D. melanogaster* genes that are not highly enriched in the ovary, according to FlyAtlas2[1]. Orange are transcripts matching genes that are highly enriched in the ovary. Black are transcripts without a BLAST match in *D. melanogaster*. Gray are non-significantly differentially expressed transcripts.

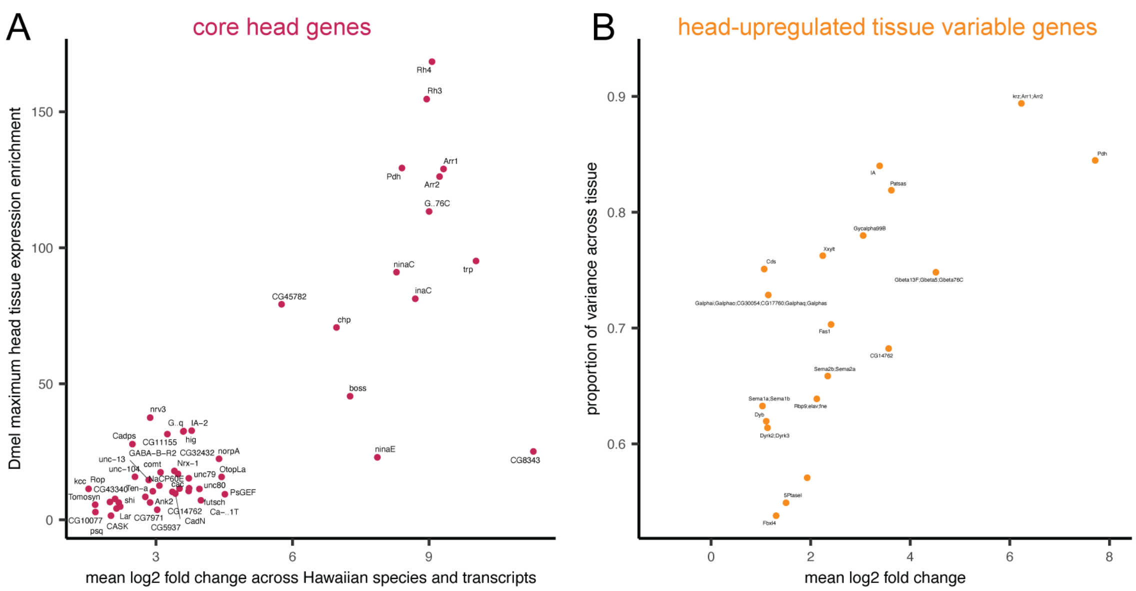

Figure S9: **Differential gene expression analyses comparing head and carcass, colored by *D. melanogaster* expression enrichment.** A, Core head genes, plotted by mean expression change across Hawaiian species to the maximum head, brain, or eye enrichment values from *D. melanogaster*, as reported in FlyAtlas2[1]. Core genes are annotated with the gene symbol from *D. melanogaster*. B, Tissue-variable genes identified in a linear model analysis comparing head and carcass tissues, showing head-upregulated TVGs. Genes are annotated with the gene symbol from the *D. melanogaster* sequences in the same homology group.

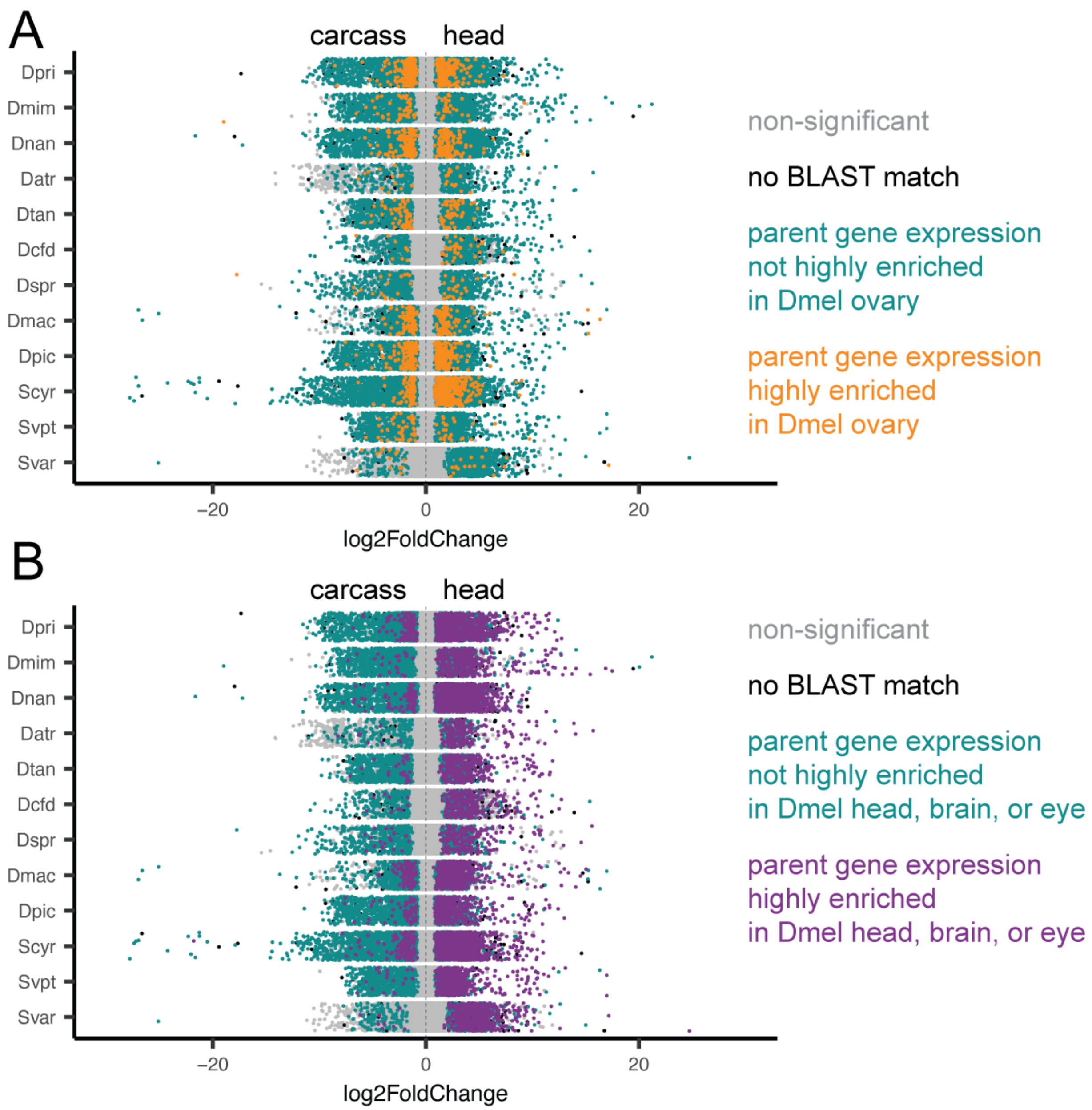

Figure S10: **Differential gene expression analyses comparing head and carcass, colored by *D. melanogaster* expression enrichment.** A, Points in cyan indicate transcripts that match *D. melanogaster* genes that are not highly enriched in the head, according to FlyAtlas2[1]. Orange are transcripts matching genes that are highly enriched in the head. Black are transcripts without a BLAST match in *D. melanogaster*. Gray are non-significantly differentially expressed transcripts. B, The same plot, with purple points as transcripts that are highly enriched in the head, brain, or eye of *D. melanogaster*.

#### 1.4 Linear modeling

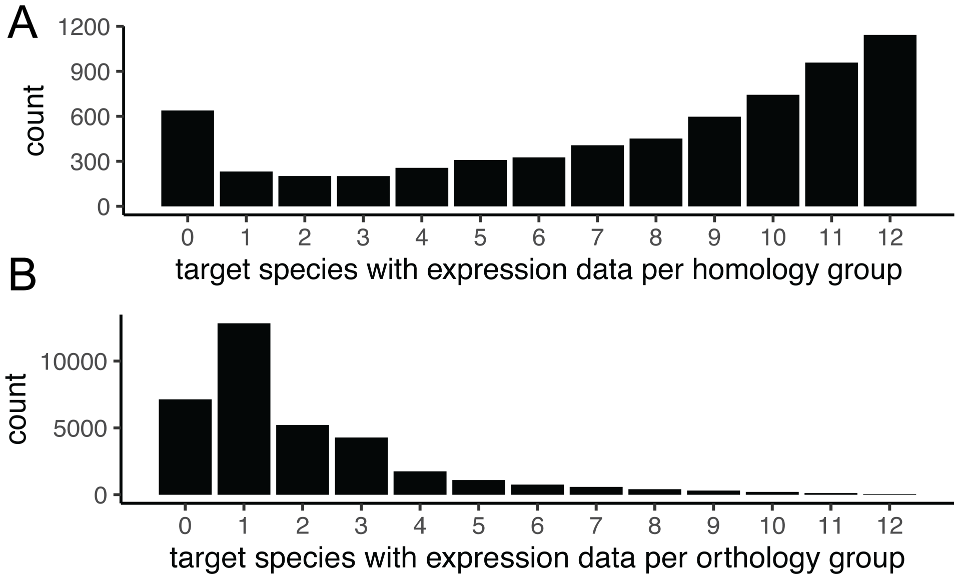

Figure S11: **Representation across homology and orthology groups.** A, The number of target species with expression data per homology group, inferred with the agalma pipeline using an all-by-all BLAST approach between the twelve reference transcriptomes and twelve other published Drosophilidae genomes. Groups with zero species are those that contain only genes from the species with published genomes, and not the twelve target species sequenced this study. B, The number of target species with expression data per orthology group, inferred with the agalma pipeline, using gene trees to identify orthologs.

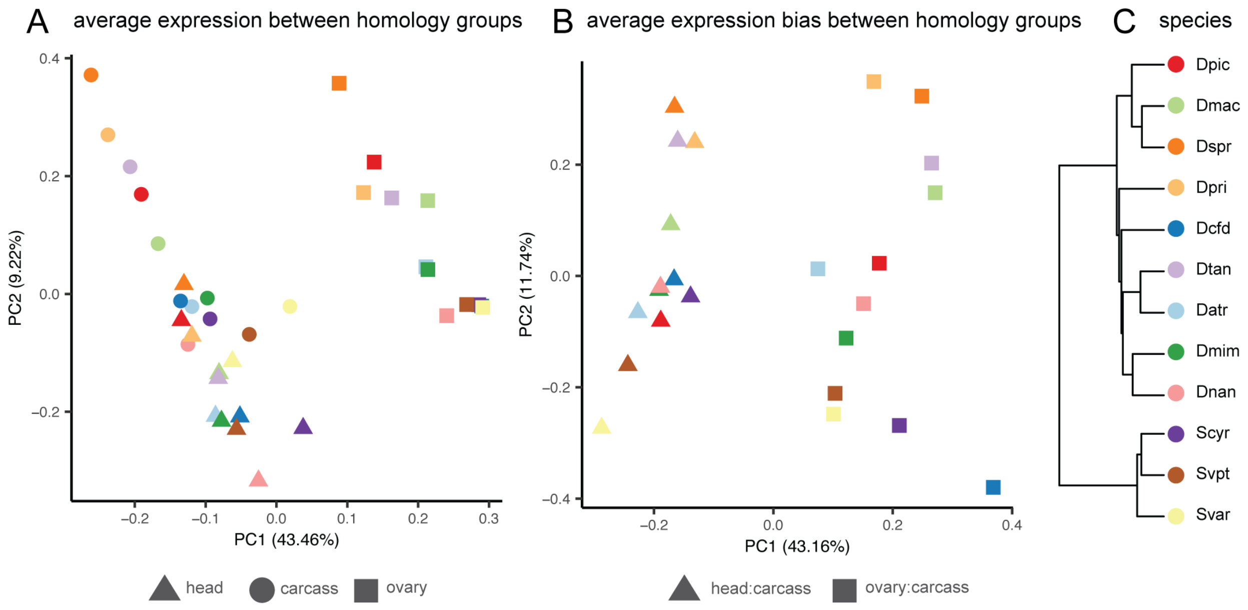

Figure S12: **Principal component analysis across species.** A, PCA using average expression across genes grouped by homology, as inferred with the agalma pipeline. B, PCA using average expression bias, calculated as an expression ratio between the ovary and carcass or head and carcass. Ratios are calculated across genes grouped by homology. C, Phylogeny of species shown in A and B.

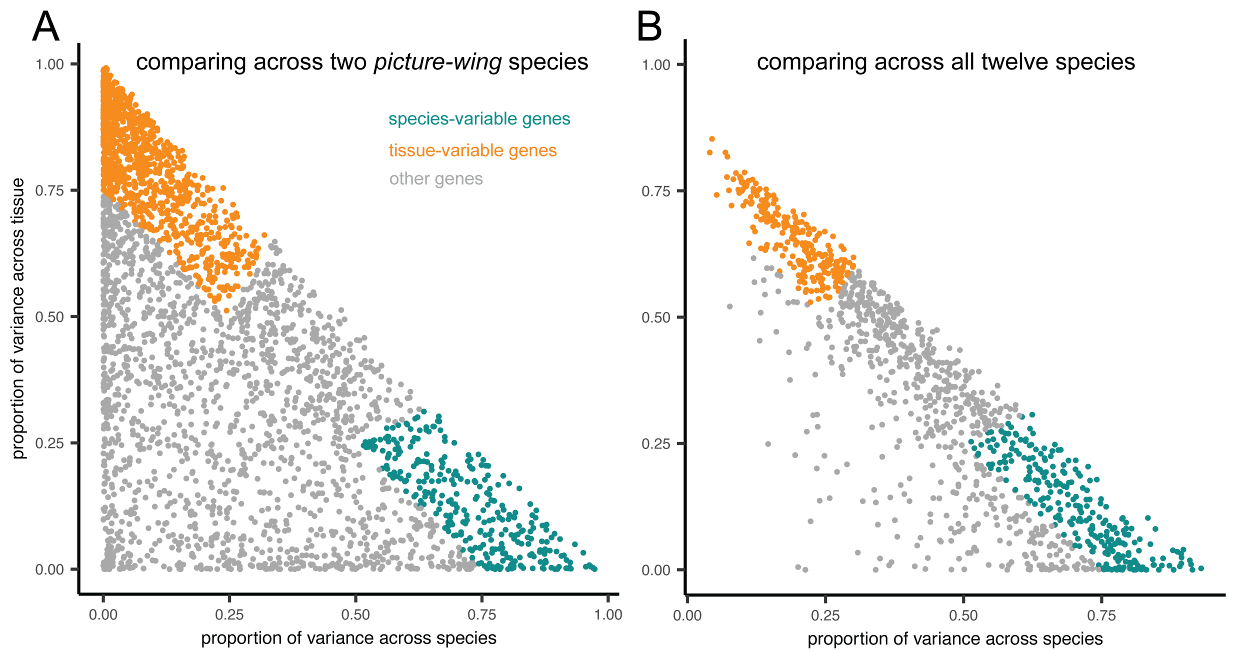

Figure S13: **Linear model results by gene.** A, Results of a linear model analysis between genes, grouped by homology as inferred in the agalma pipeline, comparing two *picture-wing* species, *D. macrothrix* and *D. sproati*. Orange points are tissue-variable genes (TVGs), cyan are species-variable genes (SVGs), gray are neither. B, Results of the same analysis across all twelve Hawaiian drosophilid species studied here.

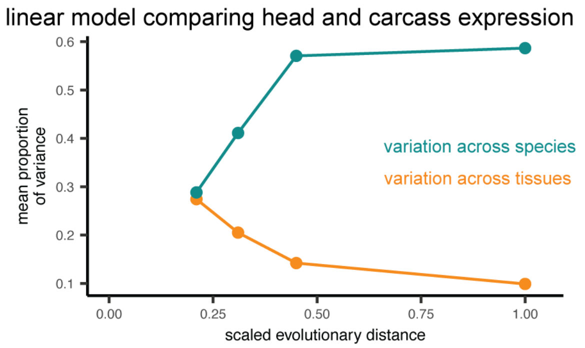

Figure S14: **Proportion of variance across species and tissues, comparing head and carcass.** The mean proportion of variance across genes attributed to species and tissues, comparing the head and carcass across four clades. Clades correspond to those shown in Fig. 3A.

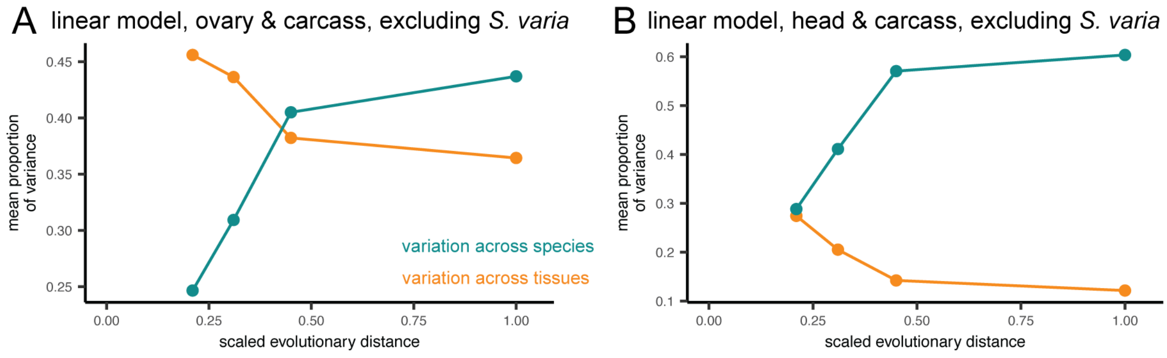

Figure S15: **Proportion of variance across species and tissues, excluding *S. varia*.** A, The mean proportion of variance across genes attributed to species and tissues, comparing the ovary and carcass across four clades, excluding *S. varia* from clade D. B, The same plot, comparing the head and carcass. Clades correspond to those shown in Fig. 3A.

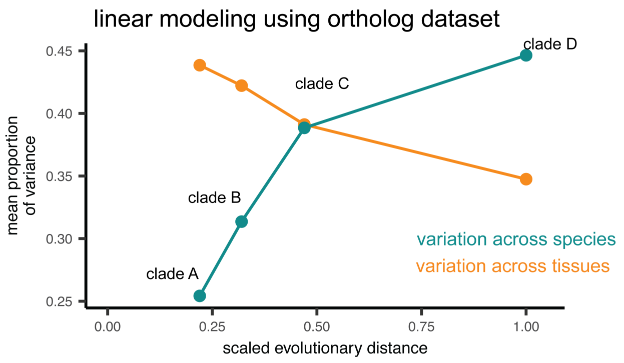

Figure S16: **Proportion of variance across species and tissues, calculated using a dataset of 1:1 orthologs.** The mean proportion of variance across genes attributed to species and tissues, comparing the ovary and carcass across four clades. Sample sizes are for clade A: 2,185 orthology groups, clade B: 918, clade C: 87, and clade D: 37.

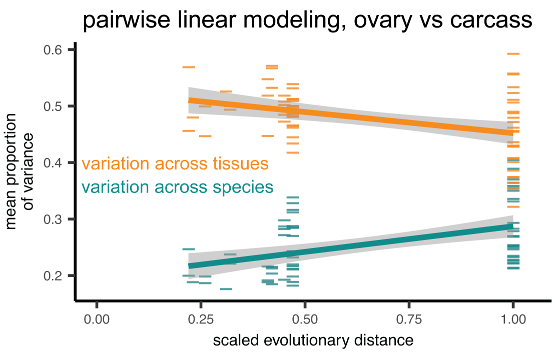

Figure S17: **Proportion of variance across species and tissues, calculated using pairwise combinations of species.** The mean proportion of variance across genes attributed to species and tissues, comparing the ovary and carcass across all pairwise combinations of twelve species, shown as colored bars. Lines and gray shadow shows the regression and 95% confidence interval using a linear model.

#### 1.5 Phylogenetic analysis

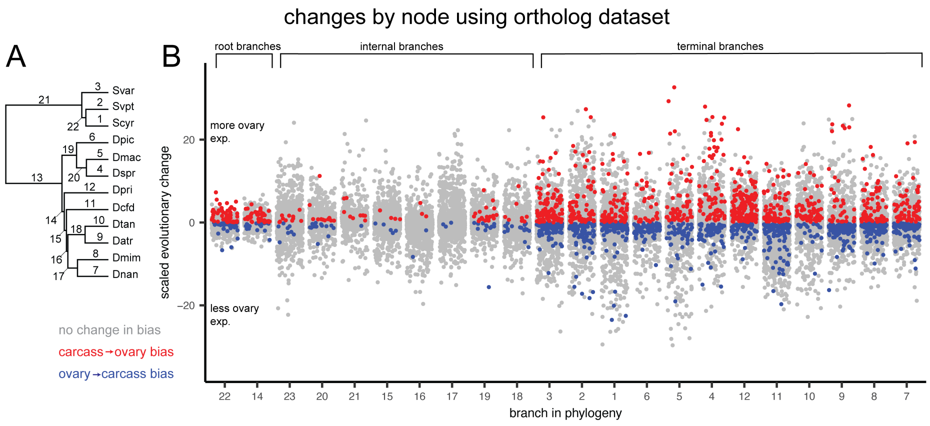

Figure S18: **Changes in ovary-biased expression in strict orthologs.** The phylogeny with all 22 branches numbered. E, The distribution of changes in expression of orthologs, grouped by branch, with random jitter on the x-axis within each group. Points colored according to the qualitative change in bias, either from more expression in ovary than carcass to less (blue), the reverse (red), or no change in overall bias (gray).

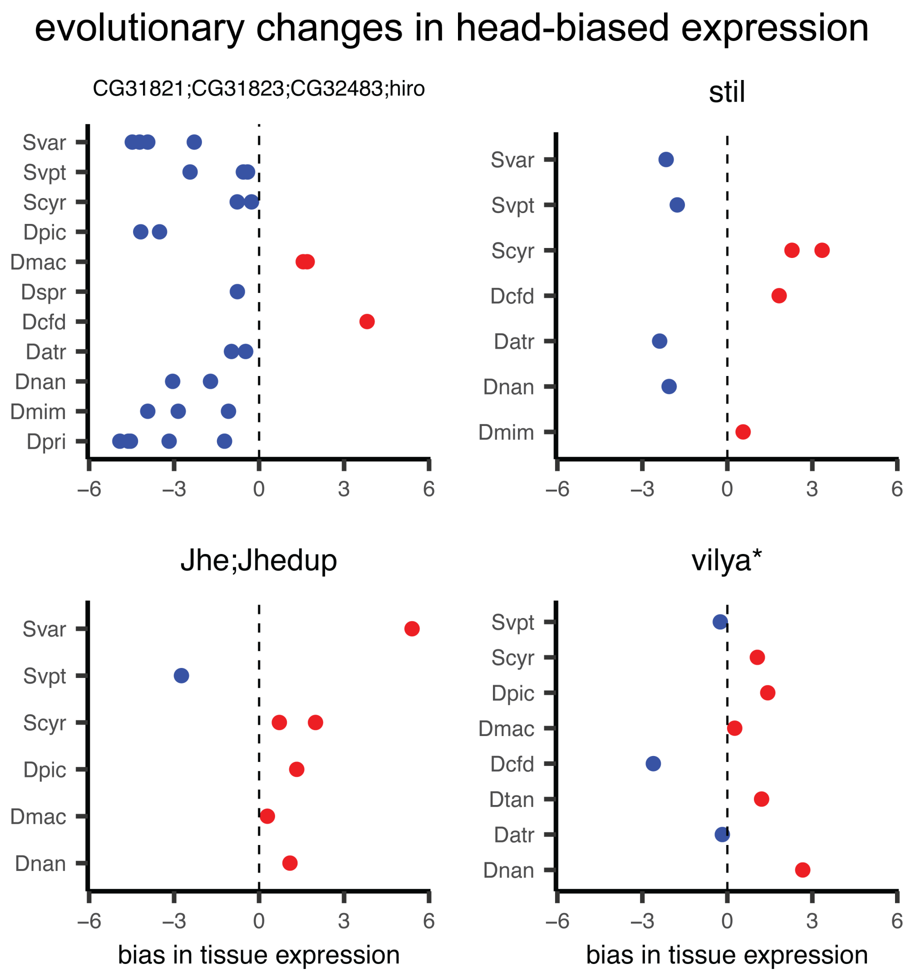

Figure S19: **Changes in head-biased expression evolution.** Four genes displaying large swings in relative head expression, showing the expression bias for each transcript colored according to more expression in the head (red) or carcass (blue). Panels are annotated with the gene symbol from the *D. melanogaster* sequences in the same homology group, with the exception of *vilya**, which was annotated using a direct BLAST search since no *D. melanogaster* sequence was present in that group.

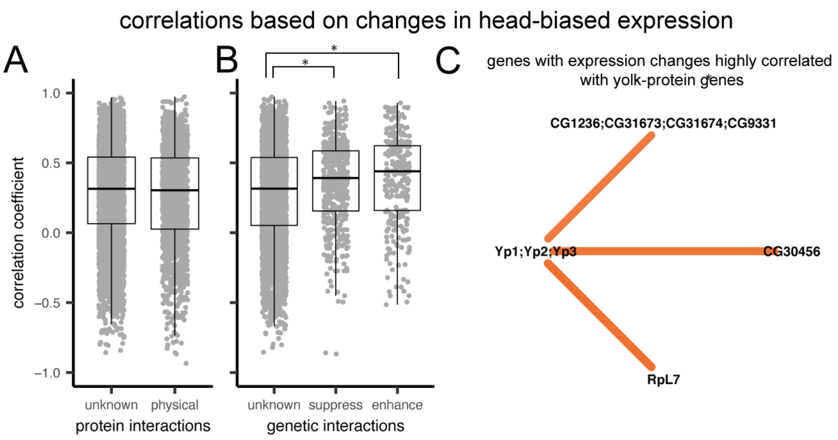

Figure S20: **Evolutionary correlations based on head-biased expression evolution.** A-B Comparison of the dsitribution of Pearson's correlation coefficients based on head-biased expression evolution between genes. Asterisks indicate a significant t-test comparison. Box plots indicate mean, upper and lower quartiles, and 1.5x interquartile ranges. A, Genes with no or unknown protein-protein interactions compared to those with reported interactions in FlyBase[2] (maximum p-value=0.256 over 100 replicates). B, Correlation comparison between genes with no or uknown genetic interactions and those reported to have enhancement or suppression interactions in FlyBase (unknown vs. enhancemenet max. p-value=<0.001; unknown vs. suppression max. p-value=<0.001; enhancement vs. suppression p-value=0.248). C, The network of strong correlation partners (absolute correlation > 0.825) with the yolk-protein genes, colored by the direction of correlation. Stronger correlations are shown by brighter colors, and thicker, shorter lines. Nodes are annotated with the gene symbols from the *D. melanogaster* sequences from that homology group.

### 2 Tables

#### 2.1 Collection and sequencing

Table S1: Field collection information for sequenced specimens.

| individual | species | general site | locality | collection method | permit | collection date | GPS |
|---|---|---|---|---|---|---|---|
| 088C | *Scaptomya cyrtandrae* | Stainback Highway | Army road west | on Cyrtandra platyphylla | DOFAW I1012; NARS special use; Hawai'i island forest reserve access permit | 5/29/2017 | N19° 33.615' W155° 15.010' |
| 088A | *Scaptomya cyrtandrae* | Stainback Highway | Army road west | on Cyrtandra platyphylla | DOFAW I1012; NARS special use; Hawai'i island forest reserve access permit | 5/29/2017 | N19° 33.615' W155° 15.010' |
| 088B | *Scaptomya cyrtandrae* | Stainback Highway | Army road west | on Cyrtandra platyphylla | DOFAW I1012; NARS special use; Hawai'i island forest reserve access permit | 5/29/2017 | N19° 33.615' W155° 15.010' |
| 025A | *Drosophila picticornis* | Koke'e State Park | Awa'awapuhi trail | baits | DOFAW I1012; Koke'e state park K2017-2015; I1012; NARS special use; Kaua'i island forest reserves KPI-2017-114 | 4/14/2017 | N22° 08.481' W159° 38.926' |
| 16.2-1 | *Drosophila cf dives* | Hawai'i Volcanoes National Park | Bird park | baits | DOFAW I1012; HAVO-2017-SCI-0017 | 5/10/2016 | N19° 26.3512' W155° 18.2225' |
| 16.1-1 | *Drosophila cf dives* | Hawai'i Volcanoes National Park | Bird park | baits | DOFAW I1012; HAVO-2017-SCI-0017 | 5/10/2016 | N19° 26.3512' W155° 18.2225' |

Table S1: Field collection information for sequenced specimens. *(continued)*

| individual | species | general site | locality | collection method | permit | collection date | GPS |
|---|---|---|---|---|---|---|---|
| 18.0-3 | *Drosophila mimica* | Hawai'i Volcanoes National Park | Bird park | sweeping Sapindus saponaria leaves | DOFAW I1012; HAVO-2017-SCI-0017 | 5/10/2016 | N19° 26.3512' W155° 18.2225' |
| 040B | *Drosophila mimica* | Hawai'i Volcanoes National Park | Bird park | sweeping Sapindus saponaria leaves | DOFAW I1012; HAVO-2017-SCI-0017 | 4/17/2017 | N19° 26.3512' W155° 18.2225' |
| 040D | *Drosophila mimica* | Hawai'i Volcanoes National Park | Bird park | sweeping Sapindus saponaria leaves | DOFAW I1012; HAVO-2017-SCI-0017 | 4/17/2017 | N19° 26.3512' W155° 18.2225' |
| 040C | *Drosophila mimica* | Hawai'i Volcanoes National Park | Bird park | sweeping Sapindus saponaria leaves | DOFAW I1012; HAVO-2017-SCI-0017 | 4/17/2017 | N19° 26.3512' W155° 18.2225' |
| 032B | *Drosophila nanella* | Koke'e State Park | Drosophila ditch | sweeping Paisonia Leaves | DOFAW I1012; Koke'e state park K2017-2015; I1012; NARS special use; Kaua'i island forest reserves KPI-2017-114 | 4/16/2017 | N22° 04.795' W159° 40.448' |
| 002C | *Drosophila nanella* | Koke'e State Park | Drosophila ditch | sweeping Paisonia Leaves | DOFAW I1012; Koke'e state park K2017-2015; I1012; NARS special use; Kaua'i island forest reserves KPI-2017-114 | 4/13/2017 | N22° 04.795' W159° 40.448' |
| 032A | *Drosophila nanella* | Koke'e State Park | Drosophila ditch | sweeping Paisonia Leaves | DOFAW I1012; Koke'e state park K2017-2015; I1012; NARS special use; Kaua'i island forest reserves KPI-2017-114 | 4/16/2017 | N22° 04.795' W159° 40.448' |
| 002D | *Drosophila nanella* | Koke'e State Park | Drosophila ditch | sweeping Paisonia Leaves | DOFAW I1012; Koke'e state park K2017-2015; I1012; NARS special use; Kaua'i island forest reserves KPI-2017-114 | 4/13/2017 | N22° 04.795' W159° 40.448' |
| 024B | *Drosophila primaeva* | Koke'e State Park | Drosophila gulch | baits | DOFAW I1012; Koke'e state park K2017-2015; I1012; NARS special use; Kaua'i island forest reserves KPI-2017-114 | 4/16/2017 | N22° 08.914' W159° 38.175' |
| 021A | *Drosophila atroscutellata* | Koke'e State Park | Nualolo trail | sweeping Corynocarpus sp leaves | DOFAW I1012; Koke'e state park K2017-2015; I1012; NARS special use; Kaua'i island forest reserves KPI-2017-114 | 4/15/2017 | N22° 07.801' W159° 39.617' |
| 029D | *Drosophila atroscutellata* | Koke'e State Park | Nualolo trail | sweeping Corynocarpus sp leaves | DOFAW I1012; Koke'e state park K2017-2015; I1012; NARS special use; Kaua'i island forest reserves KPI-2017-114 | 4/16/2017 | N22° 07.801' W159° 39.617' |
| 029A | *Drosophila atroscutellata* | Koke'e State Park | Nualolo trail | sweeping Corynocarpus sp leaves | DOFAW I1012; Koke'e state park K2017-2015; I1012; NARS special use; Kaua'i island forest reserves KPI-2017-114 | 4/16/2017 | N22° 07.801' W159° 39.617' |
| 016B | *Drosophila picticornis* | Koke'e State Park | Nualolo trail | baits | DOFAW I1012; Koke'e state park K2017-2015; I1012; NARS special use; Kaua'i island forest reserves KPI-2017-114 | 4/15/2017 | N22° 07.801' W159° 39.617' |
| 016C | *Drosophila picticornis* | Koke'e State Park | Nualolo trail | baits | DOFAW I1012; Koke'e state park K2017-2015; I1012; NARS special use; Kaua'i island forest reserves KPI-2017-114 | 4/15/2017 | N22° 07.801' W159° 39.617' |
| 028A | *Drosophila picticornis* | Koke'e State Park | Nualolo trail | baits | DOFAW I1012; Koke'e state park K2017-2015; I1012; NARS special use; Kaua'i island forest reserves KPI-2017-114 | 4/15/2017 | N22° 07.801' W159° 39.617' |
| 020B | *Scaptomyza varipicta* | Koke'e State Park | Nualolo trail | sweeping Cheirodendron sp leaves | DOFAW I1012; Koke'e state park K2017-2015; I1012; NARS special use; Kaua'i island forest reserves KPI-2017-114 | 4/15/2017 | N22° 08.336' W159° 40.479' |

Table S1: Field collection information for sequenced specimens. *(continued)*

| individual | species | general site | locality | collection method | permit | collection date | GPS |
|---|---|---|---|---|---|---|---|
| 020C | *Scaptomyza varipicta* | Koke'e State Park | Nualolo trail | sweeping Cheiroden-dron sp leaves | DOFAW I1012; Koke'e state park K2017-2015; I1012; NARS special use; Kaua'i island forest reserves KPI-2017-114 | 4/15/2017 | N22° 08.336' W159° 40.479' |
| 020D | *Scaptomyza varipicta* | Koke'e State Park | Nualolo trail | sweeping Cheiroden-dron sp leaves | DOFAW I1012; Koke'e state park K2017-2015; I1012; NARS special use; Kaua'i island forest reserves KPI-2017-114 | 4/15/2017 | N22° 08.336' W159° 40.479' |
| 020A | *Scaptomyza varipicta* | Koke'e State Park | Nualolo trail | sweeping Cheiroden-dron sp leaves | DOFAW I1012; Koke'e state park K2017-2015; I1012; NARS special use; Kaua'i island forest reserves KPI-2017-114 | 4/15/2017 | N22° 08.336' W159° 40.479' |
| 8.0-1 | *Drosophila macrothrix* | Hawai'i Volcanoes National Park | Ola'a tract, pole 44 | baits | DOFAW I1012; HAVO-2017-SCI-0017 | 5/9/2016 | N19° 27.722' W155° 14.875' |
| 8.0-2 | *Drosophila macrothrix* | Hawai'i Volcanoes National Park | Ola'a tract, pole 44 | baits | DOFAW I1012; HAVO-2017-SCI-0017 | 5/9/2016 | N19° 27.722' W155° 14.875' |
| 8.0-3 | *Drosophila macrothrix* | Hawai'i Volcanoes National Park | Ola'a tract, pole 44 | baits | DOFAW I1012; HAVO-2017-SCI-0017 | 5/9/2016 | N19° 27.722' W155° 14.875' |
| 055A | *Drosophila macrothrix* | Hawai'i Volcanoes National Park | Ola'a tract, pole 44 | baits | DOFAW I1012; HAVO-2017-SCI-0017 | 4/17/2017 | N19° 27.722' W155° 14.875' |
| 7.1-1 | *Drosophila sproati* | Hawai'i Volcanoes National Park | Ola'a tract, pole 44 | baits | DOFAW I1012; HAVO-2017-SCI-0017 | 5/9/2016 | N19° 27.722' W155° 14.875' |
| 7.2-1 | *Drosophila sproati* | Hawai'i Volcanoes National Park | Ola'a tract, pole 44 | baits | DOFAW I1012; HAVO-2017-SCI-0017 | 5/9/2016 | N19° 27.722' W155° 14.875' |
| 106B | *Drosophila sproati* | Hawai'i Volcanoes National Park | Ola'a tract, pole 44 | baits | DOFAW I1012; HAVO-2017-SCI-0017 | 5/29/2017 | N19° 27.722' W155° 14.875' |
| 106A | *Drosophila sproati* | Hawai'i Volcanoes National Park | Ola'a tract, pole 44 | baits | DOFAW I1012; HAVO-2017-SCI-0017 | 5/29/2017 | N19° 27.722' W155° 14.875' |
| 043C | *Drosophila tanythrix* | Hawai'i Volcanoes National Park | Ola'a tract, pole 44 | baits | DOFAW I1012; HAVO-2017-SCI-0017 | 4/18/2017 | N19° 27.722' W155° 14.875' |
| 056A | *Drosophila tanythrix* | Hawai'i Volcanoes National Park | Ola'a tract, pole 44 | baits | DOFAW I1012; HAVO-2017-SCI-0017 | 4/17/2017 | N19° 27.722' W155° 14.875' |
| 40.2-1 | *Drosophila tanythrix* | Hawai'i Volcanoes National Park | Ola'a tract, pole 44 | baits | DOFAW I1012; HAVO-2017-SCI-0017 | 5/9/2016 | N19° 27.722' W155° 14.875' |
| 043D | *Drosophila tanythrix* | Hawai'i Volcanoes National Park | Ola'a tract, pole 44 | baits | DOFAW I1012; HAVO-2017-SCI-0017 | 4/18/2017 | N19° 27.722' W155° 14.875' |
| 008A | *Drosophila primaeva* | Koke'e State Park | Pihea trail | baits | DOFAW I1012; Koke'e state park K2017-2015; I1012; NARS special use; Kaua'i island forest reserves KPI-2017-114 | 4/14/2017 | N22° 08.799' W159° 37.074' |
| 012A | *Drosophila primaeva* | Koke'e State Park | Pihea trail | baits | DOFAW I1012; Koke'e state park K2017-2015; I1012; NARS special use; Kaua'i island forest reserves KPI-2017-114 | 4/14/2017 | N22° 08.799' W159° 37.074' |

Table S1: Field collection information for sequenced specimens. *(continued)*

| individual | species | general site | locality | collection method | permit | collection date | GPS |
|---|---|---|---|---|---|---|---|
| 008D | *Drosophila primaeva* | Koke'e State Park | Pihea trail | baits | DOFAW I1012; Koke'e state park K2017-2015; I1012; NARS special use; Kaua'i island forest reserves KPI-2017-114 | 4/14/2017 | N22° 08.799' W159° 37.074' |
| CFC | *Scaptomya varia* | Koke'e State Park | Pihea trail | collected rotting Clermontia sp flowers | DOFAW I1012; Koke'e state park K2017-2015; I1012; NARS special use; Kaua'i island forest reserves KPI-2017-114 | 4/14/2017 | N22° 08.799' W159° 37.074' |
| CFA | *Scaptomya varia* | Koke'e State Park | Pihea trail | collected rotting Clermontia sp flowers | DOFAW I1012; Koke'e state park K2017-2015; I1012; NARS special use; Kaua'i island forest reserves KPI-2017-114 | 4/14/2017 | N22° 08.799' W159° 37.074' |
| CFB | *Scaptomya varia* | Koke'e State Park | Pihea trail | collected rotting Clermontia sp flowers | DOFAW I1012; Koke'e state park K2017-2015; I1012; NARS special use; Kaua'i island forest reserves KPI-2017-114 | 4/14/2017 | N22° 08.799' W159° 37.074' |

Table S2: DNA barcoding for identification of females.

| individual | sample | species match | reference male | external reference sequence | barcode sequence used for final identification | notes |
|---|---|---|---|---|---|---|
| 040B | 040Bb | *D. mimica* | yes | yes | 16S | |
| 040C | 040Ctxt | *D. mimica* | yes | yes | 16S | |
| 040D | 040Db | *D. mimica* | yes | yes | 16S | |
| 18.0-3 | 18.0.17 | *D. mimica* | yes | yes | 16S | |
| CFA | CFAb | *S. varia* | yes | yes | COI | |
| CFB | CFBb | *S. varia* | yes | yes | COI | |
| CFC | CFCb | *S. varia* | yes | yes | COI | |
| 088A | 088Ab | *S. cyrtandrae* | none | yes | COII | |
| 088B | 088Bb | *S. cyrtandrae* | none | yes | COII | |
| 088C | 088Cb | *S. cyrtandrae* | none | yes | COII | |
| 021A | 021Ab | *D. atroscutellata* | yes | yes | COII | |
| 029A | 029Atxt | *D. atroscutellata* | yes | yes | COII | |
| 029D | 029Db | *D. atroscutellata* | yes | yes | COII | |
| 002C | 002Cb | *D. nanella* | yes | yes | 16S, COII | |
| 002D | 002Dtxt | *D. nanella* | yes | yes | 16S, COII | |
| 032A | 032Ab | *D. nanella* | yes | yes | 16S, COII | |
| 032B | 032Bb | *D. nanella* | yes | yes | 16S, COII | |
| 106A | 106Atxt | *D. sproati* | none | none | COII | matched to other females, morphology is distinctive for females in this species |
| 106B | 106Bb | *D. sproati* | none | none | COII | matched to other females, morphology is distinctive for females in this species |
| 7.1-1 | 7.1.4 | *D. sproati* | none | none | COII | matched to other females, morphology is distinctive for females in this species |
| 043C | 043Cb | *D. tanythrix* | yes | yes | COII | barcode sequences for D. cognata and D. yooni males suggest hidden complexity in this group |
| 043D | 043Dtxt | *D. tanythrix* | yes | yes | COII | barcode sequences for D. cognata and D. yooni males suggest hidden complexity in this group |
| 056A | 056Ab | *D. tanythrix* | yes | yes | COII | barcode sequences for D. cognata and D. yooni males suggest hidden complexity in this group |
| 40.2-1 | 40.2.2 | *D. tanythrix* | yes | yes | COII | barcode sequences for D. cognata and D. yooni males suggest hidden complexity in this group |
| 020A | 020Atxt | *S. varipicta* | yes | yes | COII | |
| 020B | 020Bb | *S. varipicta* | yes | yes | COII | |
| 020C | 020Cb | *S. varipicta* | yes | yes | COII | |
| 020D | 020Db | *S. varipicta* | yes | yes | COII | |

Table S2: DNA barcoding for identification of females. *(continued)*

| individual | sample | species match | reference male | external reference sequence | barcode sequence used for final identification | notes |
|---|---|---|---|---|---|---|
| 16.1-1 | 16.1.4 | *D. cf dives* | none | none | 16S, COII | found no matching reference sequence and no males were caught |
| 16.2-1 | 16.2.4 | *D. cf dives* | none | none | 16S, COII | found no matching reference sequence and no males were caught |

Table S3: Paired-end sequencing read counts.

| species | individual ID | sample ID | tissue | reads - round 1 | reads - round 2 | reads - round 3 | total reads |
|---|---|---|---|---|---|---|---|
| *D. atroscutellata* | 029A | 029Atxt | whole fly | 12,520,922 | 7,801,177 | 19,925,042 | 40,247,141 |
| *D. cf dives* | 16.1-1 | 16.1.1 | ovary | 16,808,131 | | | 16,808,131 |
| *D. cf dives* | 16.1-1 | 16.1.2 | head | 18,215,227 | | | 18,215,227 |
| *D. cf dives* | 16.1-1 | 16.1.4 | carcass | | 8,773,601 | | 8,773,601 |
| *D. macrothrix* | 055A | 055Atxt | whole fly | 10,336,762 | 10,313,394 | 19,643,740 | 40,293,896 |
| *D. macrothrix* | 8.0-2 | 8.0.6 | ovary | 8,886,609 | | | 8,886,609 |
| *D. mimica* | 040C | 040Ctxt | whole fly | 9,471,290 | 8,887,955 | 15,751,511 | 34,110,756 |
| *D. nanella* | 002D | 002Dtxt | whole fly | 10,833,205 | 7,350,705 | 17,211,801 | 35,395,711 |
| *D. picticornis* | 025A | 025Atxt | whole fly | 10,085,602 | 10,177,523 | 16,172,455 | 36,435,580 |
| *D. primaeva* | 008D | 008Dtxt | whole fly | 9,129,075 | 7,583,577 | 13,937,132 | 30,649,784 |
| *D. sproati* | 106A | 106Atxt | whole fly | 7,432,370 | 6,026,569 | 10,237,744 | 23,696,683 |
| *D. tanythrix* | 043D | 043Dtxt | whole fly | 11,293,054 | 8,833,878 | 15,667,639 | 35,794,571 |
| *S. cyrtandrae* | 088B | 088Bb | carcass | | 9,166,453 | | 9,166,453 |
| *S. cyrtandrae* | 088B | 088Bn | head | | 8,796,864 | | 8,796,864 |
| *S. cyrtandrae* | 088B | 088Bo | ovary | | 9,763,204 | | 9,763,204 |
| *S. cyrtandrae* | 088C | 088Cb | carcass | 11,218,315 | | | 11,218,315 |
| *S. cyrtandrae* | 088C | 088Cn | head | 7,670,354 | | | 7,670,354 |
| *S. cyrtandrae* | 088C | 088Co | ovary | 7,145,307 | | | 7,145,307 |
| *S. varia* | CFB | CFBb | carcass | | 6,672,052 | | 6,672,052 |
| *S. varia* | CFB | CFBn | head | | 6,311,203 | | 6,311,203 |
| *S. varia* | CFB | CFBo | ovary | | 12,672,693 | | 12,672,693 |
| *S. varia* | CFC | CFCb | carcass | 6,941,585 | | | 6,941,585 |
| *S. varia* | CFC | CFCn | head | 8,853,906 | | | 8,853,906 |
| *S. varia* | CFC | CFCo | ovary | 8,161,304 | | | 8,161,304 |
| *S. varipicta* | 020A | 020Atxt | whole fly | 7,690,349 | 8,004,757 | 14,380,332 | 30,075,438 |

Table S4: Single-end sequencing read counts.

| species | individual ID | sample ID | tissue | notes | reads - round 1 | reads - round 2 | reads - round 3 | reads - round 4 | total reads |
|---|---|---|---|---|---|---|---|---|---|
| *D. atroscutellata* | 021A | 021Ab | carcass | ovipositor removed | | | 14,147,182 | | 14,147,182 |
| *D. atroscutellata* | 021A | 021An | head | | | | 14,091,512 | | 14,091,512 |
| *D. atroscutellata* | 021A | 021Ao | ovary | | | | 17,225,774 | | 17,225,774 |
| *D. atroscutellata* | 029D | 029Db | carcass | ovipositor removed | | | | 13,664,824 | 13,664,824 |
| *D. atroscutellata* | 029D | 029Dn | head | | | | | 18,613,630 | 18,613,630 |
| *D. atroscutellata* | 029D | 029Do | ovary | no vitel-logenic eggs | | | | 19,916,293 | 19,916,293 |
| *D. cf dives* | 16.1-1 | 16.1.1 | ovary | | 12,041,362 | | | | 12,041,362 |
| *D. cf dives* | 16.1-1 | 16.1.2 | head | | 18,382,021 | | | | 18,382,021 |
| *D. cf dives* | 16.1-1 | 16.1.4 | carcass | | 21,990,829 | | | | 21,990,829 |
| *D. cf dives* | 16.2-1 | 16.2.1 | ovary | | 16,508,284 | | | | 16,508,284 |
| *D. cf dives* | 16.2-1 | 16.2.2 | head | | | 11,288,812 | | | 11,288,812 |
| *D. cf dives* | 16.2-1 | 16.2.4 | carcass | | 14,384,696 | | | | 14,384,696 |
| *D. macrothrix* | 8.0-1 | 8.0.1 | ovary | | 625,317 | 337,147 | | | 962,464 |
| *D. macrothrix* | 8.0-1 | 8.0.2 | head | | 28,027,155 | 12,695,316 | | | 40,722,471 |
| *D. macrothrix* | 8.0-1 | 8.0.4 | carcass | | 18,125,412 | | | | 18,125,412 |
| *D. macrothrix* | 8.0-2 | 8.0.6 | ovary | | 15,583,003 | | | | 15,583,003 |
| *D. macrothrix* | 8.0-2 | 8.0.7 | head | | | | | 12,639,355 | 12,639,355 |
| *D. macrothrix* | 8.0-2 | 8.0.9 | carcass | | 42,026,251 | | | | 42,026,251 |
| *D. macrothrix* | 8.0-3 | 8.0.12 | carcass | | 29,231,317 | | | | 29,231,317 |
| *D. macrothrix* | 8.0-3 | 8.0.15 | ovary | | 35,169,909 | | | | 35,169,909 |
| *D. macrothrix* | 8.0-3 | 8.0.16 | head | | 17,682,909 | | | | 17,682,909 |
| *D. mimica* | 040B | 040Bb | carcass | | | | | 13,288,927 | 13,288,927 |

Table S4: Single-end sequencing read counts. *(continued)*

| species | individual ID | sample ID | tissue | notes | reads - round 1 | reads - round 2 | reads - round 3 | reads - round 4 | total reads |
|---|---|---|---|---|---|---|---|---|---|
| *D. mimica* | 040B | 040Bn | head | | | | | 12,473,218 | 12,473,218 |
| *D. mimica* | 040B | 040Bo | ovary | | | | | 14,622,979 | 14,622,979 |
| *D. mimica* | 040D | 040Db | carcass | | | | 20,653,357 | | 20,653,357 |
| *D. mimica* | 040D | 040Db | head | | | | 12,923,155 | | 12,923,155 |
| *D. mimica* | 040D | 040Db | ovary | | | | 17,181,968 | | 17,181,968 |
| *D. mimica* | 18.0-3 | 18.0.14 | ovary | | | | 13,450,864 | 17,450,051 | 30,900,915 |
| *D. mimica* | 18.0-3 | 18.0.15 | head | sequencing failed | | | | | 0 |
| *D. mimica* | 18.0-3 | 18.0.17 | carcass | | 19,971,465 | | | 11,948,038 | 31,919,503 |
| *D. nanella* | 002C | 002Cb | carcass | | | 15,668,215 | | | 15,668,215 |
| *D. nanella* | 002C | 002Cn | head | | | 18,850,083 | | | 18,850,083 |
| *D. nanella* | 002C | 002Co | ovary | | | 19,143,364 | | | 19,143,364 |
| *D. nanella* | 032A | 032Ab | carcass | ovipositor removed | | | 16,985,268 | | 16,985,268 |
| *D. nanella* | 032A | 032An | head | | | | 15,262,003 | | 15,262,003 |
| *D. nanella* | 032A | 032Ao | ovary | | | | 26,566,152 | | 26,566,152 |
| *D. nanella* | 032B | 032Bb | carcass | ovipositor removed | | | | 13,166,018 | 13,166,018 |
| *D. nanella* | 032B | 032Bn | head | | | | | 7,720,333 | 7,720,333 |
| *D. nanella* | 032B | 032Bo | ovary | | | | | 10,507,831 | 10,507,831 |
| *D. picticornis* | 016B | 016Bb | carcass | | | | | 13,944,238 | 13,944,238 |
| *D. picticornis* | 016B | 016Bn | head | | | | | 12,710,348 | 12,710,348 |
| *D. picticornis* | 016B | 016Bo | ovary | | | | | 8,261,788 | 8,261,788 |
| *D. picticornis* | 016C | 016Cb | carcass | ovipositor removed | | 19,183,647 | | | 19,183,647 |
| *D. picticornis* | 016C | 016Cn | head | | | 20,597,014 | | | 20,597,014 |
| *D. picticornis* | 016C | 016Co | ovary | | | 18,242,638 | | | 18,242,638 |
| *D. picticornis* | 028A | 028Ab | carcass | ovipositor removed | | | 17,704,561 | 21,917,683 | 39,622,244 |
| *D. picticornis* | 028A | 028An | head | | | | 15,060,854 | 12,034,753 | 27,095,607 |
| *D. picticornis* | 028A | 028Ao | ovary | | | | 25,988,121 | 10,261,716 | 36,249,837 |
| *D. primaeva* | 008A | 008Ab | carcass | ovipositor removed | | | | 21,917,683 | 21,917,683 |
| *D. primaeva* | 008A | 008An | head | | | | | 12,034,753 | 12,034,753 |
| *D. primaeva* | 008A | 008Ao | ovary | | | | | 10,261,716 | 10,261,716 |
| *D. primaeva* | 012A | 012Ab | carcass | | | 16,297,595 | | | 16,297,595 |
| *D. primaeva* | 012A | 012An | head | | | 12,515,877 | | | 12,515,877 |
| *D. primaeva* | 012A | 012Ao | ovary | all oocytes were at the same stage | | 14,945,335 | | | 14,945,335 |
| *D. primaeva* | 024B | 024Bb | carcass | | | | 21,578,797 | | 21,578,797 |
| *D. primaeva* | 024B | 024Bn | head | | | | 13,850,246 | | 13,850,246 |
| *D. primaeva* | 024B | 024Bo | ovary | all oocytes were at the same stage | | | 12,001,961 | | 12,001,961 |
| *D. sproati* | 106B | 106Bb | carcass | | | 17,653,753 | | | 17,653,753 |
| *D. sproati* | 106B | 106Bn | ovary | | | 10,220,778 | | | 10,220,778 |
| *D. sproati* | 106B | 106Bo | head | | | 15,615,762 | | | 15,615,762 |
| *D. sproati* | 7.1-1 | 7.1.1 | ovary | | 9,527,229 | | | | 9,527,229 |
| *D. sproati* | 7.1-1 | 7.1.2 | head | sequencing failed | | | | | 0 |
| *D. sproati* | 7.1-1 | 7.1.4 | carcass | | 13,815,824 | | | | 13,815,824 |
| *D. sproati* | 7.2-1 | 7.2.1 | carcass | | 22,462,042 | | | | 22,462,042 |
| *D. sproati* | 7.2-1 | 7.2.2 | head | | 21,654,908 | | | | 21,654,908 |
| *D. sproati* | 7.2-1 | 7.2.4 | ovary | | 31,064,408 | | | | 31,064,408 |
| *D. tanythrix* | 043C | 043Cb | carcass | | | 11,381,085 | | | 11,381,085 |
| *D. tanythrix* | 043C | 043Cn | head | | | 9,395,340 | | | 9,395,340 |
| *D. tanythrix* | 043C | 043Co | ovary | | | 13,769,669 | | | 13,769,669 |
| *D. tanythrix* | 056A | 056Ab | carcass | | | | 20,904,414 | | 20,904,414 |
| *D. tanythrix* | 056A | 056An | head | | | | 13,824,612 | | 13,824,612 |
| *D. tanythrix* | 056A | 056Ao | ovary | | | | 17,772,798 | | 17,772,798 |
| *D. tanythrix* | 40.2-1 | 40.2.1 | ovary | | 30,223,622 | | | | 30,223,622 |
| *D. tanythrix* | 40.2-1 | 40.2.2 | head | | 27,274,868 | | 17,405,688 | 17,790,177 | 62,470,733 |
| *D. tanythrix* | 40.2-1 | 40.2.4 | carcass | | 22,021,646 | 16,772,624 | | | 38,794,270 |
| *S. cyrtandrae* | 088A | 088Ab | carcass | | | 19,038,552 | | | 19,038,552 |
| *S. cyrtandrae* | 088A | 088An | head | | | 12,179,789 | | | 12,179,789 |
| *S. cyrtandrae* | 088A | 088Ao | ovary | | | 13,439,568 | | | 13,439,568 |
| *S. cyrtandrae* | 088B | 088Bb | carcass | | | | 11,543,268 | | 11,543,268 |
| *S. cyrtandrae* | 088B | 088Bn | head | | | | 12,904,077 | | 12,904,077 |
| *S. cyrtandrae* | 088B | 088Bo | ovary | | | | 11,569,554 | | 11,569,554 |
| *S. cyrtandrae* | 088C | 088Cb | carcass | | | | | 15,301,343 | 15,301,343 |
| *S. cyrtandrae* | 088C | 088Cn | head | | | | | 8,191,593 | 8,191,593 |

Table S4: Single-end sequencing read counts. *(continued)*

| species | individual ID | sample ID | tissue | notes | reads - round 1 | reads - round 2 | reads - round 3 | reads - round 4 | total reads |
|---|---|---|---|---|---|---|---|---|---|
| *S. cyrtandrae* | 088C | 088Co | ovary | | | | | 7,158,247 | 7,158,247 |
| *S. varia* | CFA | CFAb | carcass | | | 14,094,153 | | | 14,094,153 |
| *S. varia* | CFA | CFAn | head | | | 10,655,605 | | | 10,655,605 |
| *S. varia* | CFA | CFAo | ovary | | | 10,702,817 | | | 10,702,817 |
| *S. varia* | CFB | CFBb | carcass | | | | 8,049,520 | | 8,049,520 |
| *S. varia* | CFB | CFBn | head | | | | 7,357,905 | | 7,357,905 |
| *S. varia* | CFB | CFBo | ovary | | | | 12,557,066 | | 12,557,066 |
| *S. varia* | CFC | CFCb | carcass | | | | | 11,159,366 | 11,159,366 |
| *S. varia* | CFC | CFCn | head | | | | | 11,517,456 | 11,517,456 |
| *S. varia* | CFC | CFCo | ovary | | | | | 10,874,670 | 10,874,670 |
| *S. varipicta* | 020B | 020Bb | carcass | | | 8,132,403 | | | 8,132,403 |
| *S. varipicta* | 020B | 020Bn | head | | | 14,331,603 | | | 14,331,603 |
| *S. varipicta* | 020B | 020Bo | ovary | | | 8,192,636 | | | 8,192,636 |
| *S. varipicta* | 020C | 020Cb | carcass | ovipositor removed | | | 16,584,774 | | 16,584,774 |
| *S. varipicta* | 020C | 020Cn | head | | | | 13,783,546 | | 13,783,546 |
| *S. varipicta* | 020C | 020Co | ovary | | | | 17,539,765 | | 17,539,765 |
| *S. varipicta* | 020D | 020Db | carcass | ovipositor removed | | | | 19,358,586 | 19,358,586 |
| *S. varipicta* | 020D | 020Dn | head | | | | | 19,832,422 | 19,832,422 |
| *S. varipicta* | 020D | 020Do | ovary | | | | | 13,226,137 | 13,226,137 |

Table S5: Number of transcripts per homology group per species.

| species | homology groups | groups with more than one transcript | max num. transcripts per group | ave num. transcripts per group |
|---|---|---|---|---|
| *D. atroscutellata* | 3,939 | 1,211 | 8 | 1.40 |
| *D. cf dives* | 4,782 | 966 | 8 | 1.28 |
| *D. macrothrix* | 4,061 | 1,164 | 8 | 1.39 |
| *D. mimica* | 4,278 | 1,344 | 8 | 1.42 |
| *D. nanella* | 4,790 | 1,271 | 7 | 1.36 |
| *D. picticornis* | 2,769 | 784 | 6 | 1.37 |
| *D. primaeva* | 3,274 | 935 | 7 | 1.39 |
| *D. sproati* | 3,380 | 767 | 8 | 1.30 |
| *D. tanythrix* | 3,618 | 1,015 | 7 | 1.38 |
| *S. cyrtandrae* | 4,966 | 1,283 | 7 | 1.35 |
| *S. varia* | 4,733 | 1,109 | 12 | 1.32 |
| *S. varipicta* | 4,696 | 1,195 | 8 | 1.35 |

#### 2.2 Core genes

Table S6: Core ovary genes.

| Dmel parent gene ID | Dmel parent gene symbol | transcripts in homology group | homology group Dmel sequence symbols |
|---|---|---|---|
| FBgn0250816 | AGO3 | none | none |
| FBgn0087040 | alphaTub67C | none | none |
| FBgn0266111 | ana3 | none | none |
| FBgn0028343 | Ankle2 | some | Ankle2 |
| FBgn0041164 | armi | none | none |
| FBgn0029094 | asf1 | all | asf1 |
| FBgn0005386 | ash1 | none | none |
| FBgn0000140 | asp | none | none |
| FBgn0000182 | BicC | none | none |
| FBgn0033155 | Br140 | some | Br140 |
| FBgn0010300 | brat | some | brat |
| FBgn0263855 | BubR1 | some | BubR1 |
| FBgn0039680 | Cap | none | none |
| FBgn0027512 | CG10254 | none | none |
| FBgn0037021 | CG11399 | some | CG11399 |
| FBgn0038968 | CG12499 | none | none |
| FBgn0031070 | CG12702 | none | none |
| FBgn0039640 | CG14516 | none | none |
| FBgn0034498 | CG16868 | none | none |
| FBgn0052344 | CG32344 | none | none |
| FBgn0031875 | CG3430 | some | CG3430 |
| FBgn0250754 | CG42232 | none | none |
| FBgn0029733 | CG6927 | some | CG6927;pHCl;SecCl |

Table S6: Core ovary genes. *(continued)*

| Dmel parent gene ID | Dmel parent gene symbol | transcripts in homology group | homology group Dmel sequence symbols |
|---|---|---|---|
| FBgn0034187 | CG6967 | some | CG6701;CG6967 |
| FBgn0031947 | CG7154 | some | CG7154 |
| FBgn0034073 | CG8414 | all | CG8414 |
| FBgn0037664 | CG8420 | none | none |
| FBgn0031769 | CG9135 | some | CG9135 |
| FBgn0000307 | chif | none | none |
| FBgn0044324 | Chro | none | none |
| FBgn0261016 | clos | some | no Dmel match |
| FBgn0033890 | Ctf4 | some | Ctf4 |
| FBgn0000392 | cup | none | none |
| FBgn0000404 | CycA | some | CycA |
| FBgn0000405 | CycB | some | CycB |
| FBgn0015625 | CycB3 | some | CycB3 |
| FBgn0010382 | CycE | some | CycE |
| FBgn0039016 | Dcr | some | no Dmel match |
| FBgn0000463 | Dl | none | none |
| FBgn0262619 | DNAlig1 | none | none |
| FBgn0259113 | DNApol | none | none |
| FBgn0264326 | DNApol | none | none |
| FBgn0002905 | DNApol | none | none |
| FBgn0263600 | DNApol | none | none |
| FBgn0002183 | dre4 | none | none |
| FBgn0000996 | dup | some | dup;Pms2 |
| FBgn0087008 | e(y)3 | some | e(y)3 |
| FBgn0087008 | e(y)3 | some | no Dmel match |
| FBgn0000615 | exu | some | exu |
| FBgn0033354 | FANCI | some | CG33680;FANCI |
| FBgn0005390 | fs(1)M3 | none | none |
| FBgn0004650 | fs(1)N | none | none |
| FBgn0001085 | fz | all | fz |
| FBgn0283682 | Ge | none | none |
| FBgn0015391 | glu | none | none |
| FBgn0266580 | Gp210 | none | none |
| FBgn0261278 | grp | some | grp |
| FBgn0283499 | InR | none | none |
| FBgn0011604 | Iswi | none | none |
| FBgn0015396 | jumu | some | jumu |
| FBgn0030268 | Klp10A | some | Klp10A;Klp59C;Klp59D |
| FBgn0038476 | kuk | none | none |
| FBgn0034657 | LBR | some | LBR |
| FBgn0019686 | lok | all | lok |
| FBgn0283521 | lola | some | no Dmel match |
| FBgn0283521 | lola | some | pre |
| FBgn0039972 | Marf1 | some | Marf1 |
| FBgn0025743 | mbt | some | mbt |
| FBgn0014861 | Mcm2 | some | Mcm2 |
| FBgn0284442 | Mcm3 | some | Mcm3 |
| FBgn0017577 | Mcm5 | none | none |
| FBgn0025815 | Mcm6 | none | none |
| FBgn0020633 | Mcm7 | none | none |
| FBgn0004419 | me31B | all | me31B |
| FBgn0261786 | mi | none | none |
| FBgn0036486 | Msh6 | some | Msh6 |
| FBgn0010431 | mtrm | some | no Dmel match |
| FBgn0002872 | mu2 | some | mu2 |
| FBgn0002873 | mud | none | none |
| FBgn0002878 | mus101 | none | none |
| FBgn0002924 | ncd | none | none |
| FBgn0029970 | Nek2 | some | Nek2 |
| FBgn0032848 | nesd | all | nesd |
| FBgn0002962 | nos | all | nos |
| FBgn0039559 | NSD | none | none |
| FBgn0021761 | Nup154 | none | none |
| FBgn0033766 | Nup188 | some | Nup188 |
| FBgn0031078 | Nup205 | none | none |
| FBgn0034310 | Nup75 | all | Nup75 |
| FBgn0004882 | orb | some | orb |
| FBgn0003015 | osk | none | none |
| FBgn0003028 | ovo | some | no Dmel match |
| FBgn0003044 | Pcl | none | none |
| FBgn0261811 | pico | some | pico |
| FBgn0004872 | piwi | none | none |
| FBgn0036354 | Poc1 | some | Poc1 |
| FBgn0003124 | polo | some | polo |
| FBgn0003165 | pum | some | tna |
| FBgn0261987 | Pxt | none | none |
| FBgn0003187 | qua | none | none |

Table S6: Core ovary genes. *(continued)*

| Dmel parent gene ID | Dmel parent gene symbol | transcripts in homology group | homology group Dmel sequence symbols |
|---|---|---|---|
| FBgn0003346 | RanGAP | all | RanGAP |
| FBgn0039644 | rdog | none | none |
| FBgn0264493 | rdx | some | rdx |
| FBgn0040290 | RecQ4 | some | CG6888;Jafrac1;Jafrac2;Prx3;RecQ4 |
| FBgn0020379 | Rfx | some | Rfx |
| FBgn0034249 | RhoGAP54D | some | RhoGAP54D |
| FBgn0034249 | RhoGAP54D | some | dgt3 |
| FBgn0050085 | Rif1 | some | Rif1 |
| FBgn0250850 | rig | none | none |
| FBgn0011703 | RnrL | some | RnrL |
| FBgn0003268 | rod | some | rod |
| FBgn0025802 | Sbf | none | none |
| FBgn0032475 | Sfmbt | none | none |
| FBgn0003401 | shu | some | shu |
| FBgn0051163 | SKIP | some | SKIP |
| FBgn0016070 | smg | some | CG5280;smg |
| FBgn0016070 | smg | some | no Dmel match |
| FBgn0037025 | Spc105R | none | none |
| FBgn0027500 | spd | none | none |
| FBgn0003483 | spn | none | none |
| FBgn0033348 | Spt | some | Spt;Sry |
| FBgn0262733 | Src64B | none | none |
| FBgn0002466 | sti | none | none |
| FBgn0052676 | stx | some | stx |
| FBgn0003655 | swa | all | swa |
| FBgn0031030 | Tao | none | none |
| FBgn0003701 | thr | none | none |
| FBgn0284220 | Top2 | none | none |
| FBgn0033636 | tou | none | none |
| FBgn0041775 | tral | some | tral |
| FBgn0086356 | tum | some | Echs1;RacGAP84C;tum |
| FBgn0260780 | wisp | some | wisp |
| FBgn0039338 | XNP | some | XNP |
| FBgn0039338 | XNP | some | no Dmel match |
| FBgn0004649 | yl | none | none |
| FBgn0259789 | zld | some | no Dmel match |

Table S7: Core head genes.

| Dmel parent gene ID | Dmel parent gene symbol | transcripts in homology group | homology group Dmel sequence symbols |
|---|---|---|---|
| FBgn0261788 | Ank2 | none | none |
| FBgn0000120 | Arr1 | all | Arr1;Arr2;krz |
| FBgn0000121 | Arr2 | some | Arr1;Arr2;krz |
| FBgn0000206 | boss | all | boss |
| FBgn0264386 | Ca | none | none |
| FBgn0263111 | cac | none | none |
| FBgn0015609 | CadN | none | none |
| FBgn0053653 | Cadps | none | none |
| FBgn0013759 | CASK | none | none |
| FBgn0035720 | CG10077 | none | none |
| FBgn0039927 | CG11155 | none | none |
| FBgn0033250 | CG14762 | some | CG14762 |
| FBgn0052432 | CG32432 | none | none |
| FBgn0263077 | CG43340 | none | none |
| FBgn0029834 | CG5937 | none | none |
| FBgn0035253 | CG7971 | some | CG7971 |
| FBgn0040502 | CG8343 | some | no Dmel match |
| FBgn0267435 | chp | none | none |
| FBgn0000346 | comt | none | none |
| FBgn0285925 | Fas1 | some | Fas1 |
| FBgn0259108 | futsch | none | none |
| FBgn0027575 | GABA | some | GABA |
| FBgn0004435 | Galphaq | some | CG17760;CG30054;Galphai;Galphao;Galphaq;Galphas |
| FBgn0004623 | Gbeta76C | some | Gbeta13F;Gbeta5;Gbeta76C |
| FBgn0010114 | hig | some | hig |
| FBgn0031294 | IA | some | IA |
| FBgn0004784 | inaC | none | none |
| FBgn0261794 | kcc | some | kcc |
| FBgn0000464 | Lar | none | none |
| FBgn0267429 | lovit | some | lovit;Slc45 |
| FBgn0085434 | NaCP60E | none | none |
| FBgn0002938 | ninaC | none | none |
| FBgn0002940 | ninaE | some | ninaE;Rh2;Rh6 |
| FBgn0262738 | norpA | none | none |

Table S7: Core head genes. *(continued)*

| Dmel parent gene ID | Dmel parent gene symbol | transcripts in homology group | homology group Dmel sequence symbols |
|---|---|---|---|
| FBgn0032946 | nrv3 | some | nrv1;nrv2;nrv3 |
| FBgn0038975 | Nrx | some | Spn88Ea;Spn88Eb |
| FBgn0259994 | OtopLa | some | OtopLa |
| FBgn0285944 | para | none | none |
| FBgn0011693 | Pdh | some | Pdh |
| FBgn0264598 | PsGEF | some | no Dmel match |
| FBgn0263102 | psq | some | psq |
| FBgn0003249 | Rh3 | some | Rh3;Rh4;Rh5 |
| FBgn0003250 | Rh4 | all | Rh3;Rh4;Rh5 |
| FBgn0004574 | Rop | none | none |
| FBgn0003392 | shi | none | none |
| FBgn0267001 | Ten | none | none |
| FBgn0030412 | Tomosyn | some | CSN1a;CSN1b |
| FBgn0003861 | trp | none | none |
| FBgn0025726 | unc | none | none |
| FBgn0267002 | unc | none | none |
| FBgn0038693 | unc79 | none | none |
| FBgn0039536 | unc80 | none | none |
